# 3D Mapping of Intact Ovaries Reveals the Aging Dynamics of the Ovarian Reserve

**DOI:** 10.1101/2025.11.07.686728

**Authors:** Arturo D’Angelo, Daniel Franco-Barranco, Marco Musy, James Sharpe, Ignacio Arganda-Carreras, Elvan Böke

## Abstract

Female fertility depends on a finite pool of oocytes that depletes during aging, yet the spatiotemporal dynamics of this depletion remain poorly understood. Traditional methods obscure the 3D architecture of the ovary, limiting quantitative insights.

Here, we combine light-sheet microscopy, artificial intelligence (AI)-driven segmentation, and mathematical modeling to map over 85,000 oocytes in whole-ovaries across the reproductive lifespan in mouse. We find that newly activated oocytes represent a fixed fraction of the ovarian reserve, despite an age-related decline in total oocyte numbers. Spatial analysis revealed that oocytes are enriched along the lateral ovarian axis, and local oocyte density positively correlates with activation. We also uncover a bimodal distribution of oocyte sizes, suggesting a bottleneck during oogenesis. Finally, a differential equation-based model captures the kinetics of oocyte activation and loss. Our findings establish a quantitative framework for understanding ovarian aging and suggest that an organ-scale regulatory mechanism coordinates the age-related decline in oocyte numbers.

## Introduction

Reproductive lifespan in most mammals is defined by a finite supply of immature egg cells, termed oocytes, which are established during fetal life and progressively depleted with age. In humans, females are born with approximately 1–2 million oocytes, which decrease to ∼400,000 by puberty, and fewer than 1,000 remain by the time of menopause^1-3^. Despite this initial abundance, only about 400 oocytes are ovulated during a woman’s reproductive life, with the remainder lost through cell death. Yet, the biological mechanisms that govern the rate and pattern of this decline remain poorly understood.

Oocytes reside in specialized structures called ovarian follicles, which consist of the oocyte and surrounding somatic cells. Primordial oocytes are enclosed in primordial follicles and are considered dormant as they do not grow or divide^3^. Once activated, follicles progress through primary, secondary, and antral stages, culminating in ovulation of a fertilizable egg^4^.

The dynamics of follicle activation and depletion underpin key transitions in reproductive aging, and dysregulation of these processes contributes to disorders such as primary ovarian insufficiency and premature menopause^5-7^. Most of our current understanding of follicle dynamics comes from two-dimensional (2D) histological analyses, which provide high-resolution snapshots of ovarian architecture but sacrifice volumetric context. As a result, how oocytes are distributed in three-dimensional (3D) space and how this spatial organization changes with age remain largely unknown. Similarly, the relationship between follicular location, growth stage, and activation dynamics is still not fully elucidated. To address these limitations in a mammalian system, we present a comprehensive spatial and temporal map of the ovarian reserve using whole-organ imaging and quantitative analysis. By combining high-resolution light-sheet microscopy, AI-assisted image segmentation, and mathematical modeling, we quantified every oocyte in whole mouse ovaries across the reproductive lifespan. This approach allowed us to investigate how oocyte number, size, and spatial distribution change with age, and to uncover key principles of follicle activation dynamics.

## Results

### Whole-ovary imaging and automated oocyte quantification

The mouse has long served as a fundamental model organism in reproductive biology due to its genetic tractability, well-characterized ovarian physiology, and similarity to mammalian ovarian development and function^1,8^. Importantly, house mice *(Mus musculus)* also undergo significant changes in their ovarian reserve that parallel fluctuations observed in humans^9^, and exhibit age-related declines in fertility and ultimately cease ovulation with age^1,8^. The availability of inbred mouse colonies provides a unique opportunity to map ovarian reserve numbers by age in genetically identical large populations. We therefore employed the inbred C57BL/6J (BL6) strain to investigate ovarian reserve changes during aging.

A key challenge in quantifying ovarian reserve lies in the ovary’s complex anatomy. The mouse ovary is relatively large, measuring several millimeters in diameter, and has a non-uniform shape, which poses significant limitations for imaging due to restricted penetration depth and limited field of view. Historically, quantifying oocyte numbers in adult mouse ovaries has largely relied on manual counting from serial histological sections and subsequent normalization^9^. While widely used, this approach is labor-intensive, time-consuming, and sacrifices the three-dimensional spatial information on the oocyte’s position within the ovarian tissue. Moreover, section-based methods introduce inaccuracies in estimating primordial oocyte numbers^10^.

Recent advances in light-sheet microscopy offer an advantage over these section-based methods to image whole adult ovaries^11,12^. To overcome these limitations and enable whole-organ, high-resolution imaging of the whole ovaries, we used selective plane illumination microscopy (SPIM) with tissue clearing^13^.

Specifically, we optimized both immunostaining and clearing protocols to achieve the identification of all oocytes within the ovary: we used a commercially available DDX4 antibody to label oocytes and applied TO-PRO-3 nuclear dye to counterstain all nuclei in ovaries isolated from 5-week-old mice (Fig. 1A–D, Movie S1). Our optimized protocol enabled identification of all oocytes *in situ*. It also allowed us to distinguish oocytes at different developmental stages based on their size (Fig. 1D), providing a powerful platform to investigate ovarian architecture and follicular composition in their native spatial context (Fig. 1B–C).

**Figure 1.**
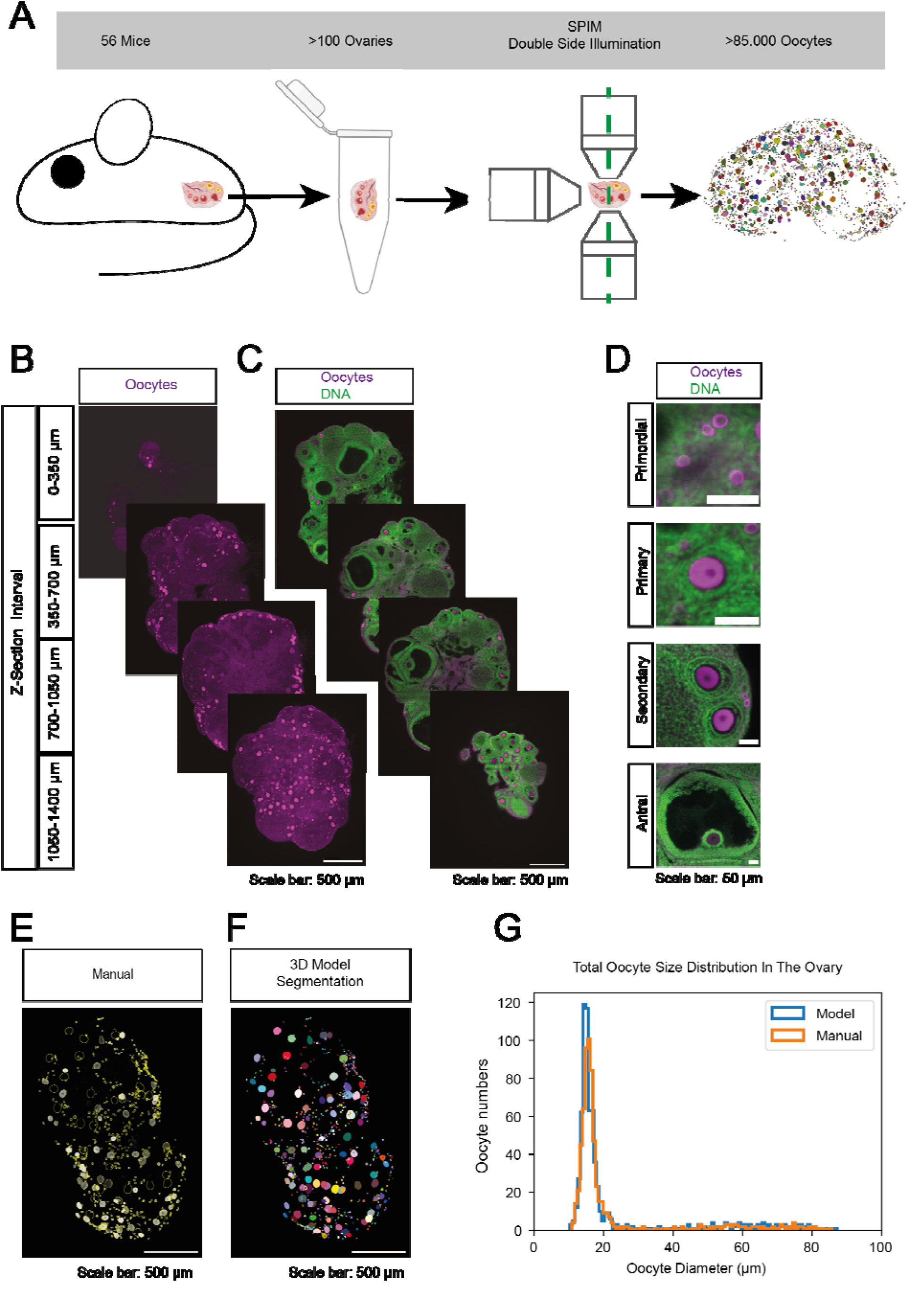
Light-sheet imaging and automated analysis of oocyte size in adult mouse Ovaries. **(A)** Cartoon summarizing the dataset size and the analysis pipeline. **(B)** Light-sheet (SPIM) imaging of whole-mount ovaries from 10-week-old C57BL/6J mice labeled with an antibody against DDX4, an oocyte marker. Images are shown as maximum-intensity projections across four z-sections, as indicated on the left, for improved visualization. **(C)** Single-plane images from 10-week-old C57BL/6J mice co-labeled with an anti-DDX4 antibody to mark oocytes and TO-PRO-3 dye to mark DNA. **(D)** Enlarged regions of interest (ROIs) from (C) highlighting different stages of folliculogenesis. **(E)** Maximum-intensity projected SPIM images of whole-mount ovaries from 5-week-old C57BL/6J mice labeled with DDX4 and manually segmented (yellow). **(F)** Same ovary as in (E), segmented using the 3D model. **(G)** Comparison of total ovarian reserve quantification between manual segmentation (E) and model-based segmentation (F).

We first applied our optimized imaging and analysis pipeline to whole ovaries from 5-week-old mice to quantify their ovarian reserve. In the inbred BL6 strain we used, females of this age are considered to be in the pubertal to early postpubertal period^14^ therefore, we selected 5 weeks as the time point to assess the maximum ovarian reserve following the first wave of oocyte activation and death^15^.

To this end, we manually segmented oocytes at different stages of activation from 3D-reconstructed SPIM image volumes, generating a curated dataset for training a deep learning model within the BiaPy framework^16^. This instance segmentation model was designed to automatically identify and delineate individual oocytes in cleared ovarian tissue (Fig. S1A). To assess its performance, we imaged five independent ovaries and compared the AI-generated segmentations with expert annotations, both before and after minimal user-guided correction (Fig. 1E–F). The model achieved a 95.3% agreement with manual segmentation, which increased to 96.7% following correction, highlighting its robustness and suitability for fast and accurate oocyte quantification (Fig. S1B–C).

This automated approach enabled us to reconstruct the full oocyte size distribution in adult mouse ovaries (Fig. 1G, Movie S2) and, notably, proved effective for quantifying oocytes in cleared human ovarian cortex samples as well (Fig. S2A–C, Movie S3), highlighting the pipeline’s cross-species applicability and its potential utility in translational and clinical research settings.

Together, these results demonstrate a robust and scalable platform to enable spatially resolved, high-throughput quantification of the ovarian reserve in the intact ovarian tissue.

### Total oocyte quantification reveals high heterogeneity between isogenic animals

Anti-Müllerian hormone (AMH), a clinical marker of ovarian reserve, exhibits substantial variability among humans, raising the possibility that oocyte numbers may also differ significantly between animals, even within genetically identical populations. To directly address this possibility, we analyzed eight 5-week-old BL6 females and quantified the total number of oocytes per animal using our whole-ovary imaging and segmentation pipeline.

Surprisingly, we observed up to a threefold variation in total oocyte number among same-aged individuals with identical genetic backgrounds, revealing substantial inter-individual variability in oocyte numbers in isogenic mice (Fig. S3A). Notably, this variability was already present in 5-week-old animals, suggesting that it arises before sexual maturation.

Given that asymmetrical ovarian responses have been reported in women undergoing hormonal stimulation^17^, we asked whether a similar left–right asymmetry exists under normal physiological conditions. To this end, we separated the left and right ovaries from each animal and quantified the ovarian reserve in each independently, but we found little asymmetry in total oocyte count between left and right ovaries in 5-week-old animals (Fig. S3B).

These findings reveal unexpectedly high variability in ovarian reserve among genetically identical animals, emphasizing the need to consider individual differences when assessing the ovarian reserve, even from genetically identical populations.

### Oocyte reserve declines with age, but transitioning fraction remains stable

Oocytes are classified according to their size and the surrounding follicular structure^1,18,19^. To investigate oocyte growth kinetics, we used the segmentation masks generated by our AI model to calculate the oocyte diameters and plot their distribution in 5-week-old ovaries. This analysis revealed a sharp peak around 15_µm, followed by a long tail corresponding to larger oocytes (Fig. 1G).

To better characterize the oocyte population around the 15 µm peak, we performed high-resolution confocal imaging of oocytes between 12–40 µm in diameter. Within this range, we could clearly distinguish dormant primordial follicles, identified by flattened squamous granulosa cells^20^, and growing primary follicles, which contain cuboidal granulosa cells (Fig. S3C–D). We also identified transitioning follicles, characterized by mixed squamous and cuboidal cell morphologies^21^. Based on size and morphology, we classified oocytes into primordial (<16 µm), transitioning (16–20 µm), and primary (20–40 µm) stages (Fig. S3C–D), enabling distinction between dormant and recently activated populations.

Next, we investigated how the ovarian reserve changes with age. Given the substantial inter-animal variability (Fig. 2B), we analyzed 56 virgin mice spanning 5 to 60 weeks of age, using at least five mice per age. Employing our high-throughput AI pipeline, we segmented over 85,000 oocytes from more than 100 intact 3D-imaged ovaries (Fig. 2A–B). This large dataset revealed substantial inter-individual variability in total oocyte numbers, with relatively lower intra-individual differences between the left and right ovaries of the same animal across all age groups (Fig. S4A–C), confirming that the high heterogeneity observed at 5 weeks persists throughout aging (Fig. S3A). As expected, total oocyte numbers declined significantly with age (Fig. 2B). The absolute numbers of primordial, transitioning, and growing oocytes all decreased as the ovarian reserve diminished. However, their relative contributions to the total population shifted: primordial oocytes decreased from ∼70% to ∼55% of the pool, while growing oocytes displayed the opposite trend, increasing from ∼15% to ∼30% (Fig. 2C).

**Figure 2.**
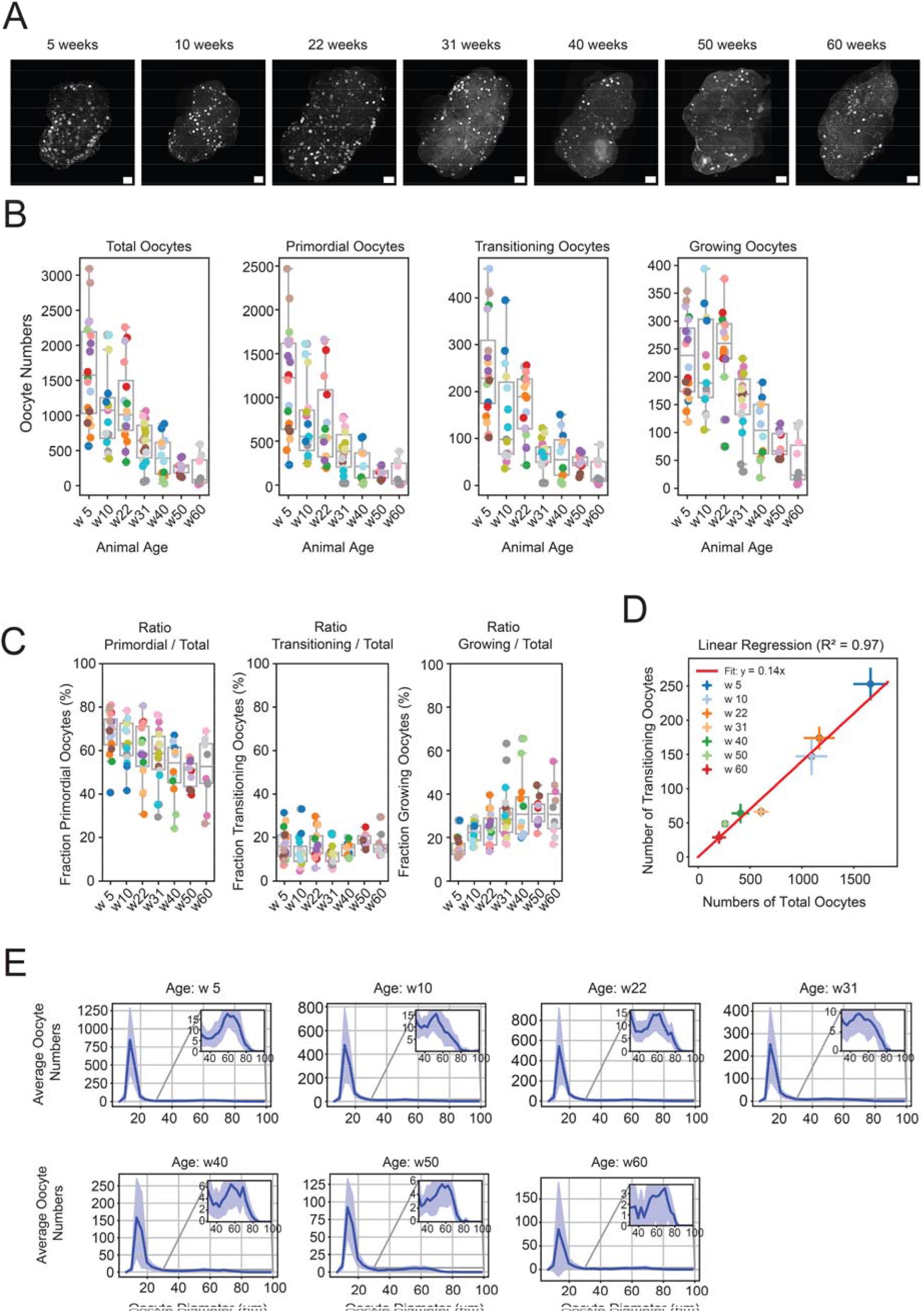
Ovarian reserve declines with age, but transitioning oocyte fraction remains stable. **(A)** Maximum-intensity projected SPIM images of ovaries from 5-, 10-, 22-, 31-, 40-, 50-, and 60-week-old inbred C57BL/6J mice labeled with an antibody against DDX4, an oocyte marker. The scale bar is 200 μm. **(B)** Quantification of the number of total, primordial, transitioning and growing oocytes. Each dot represents an ovary. Ovaries from the same animal share the same color. **(C)** Number of primordial oocytes divided by the number of total oocytes (left), number of transitioning oocytes divided by the number of total oocytes (center) and number of growing oocytes divided by the number of total oocytes (right). Each dot represents an ovary. Ovaries from the same animal share the same color. **(D)** Quantification of transitioning oocyte number as a function of total oocyte number color-coded by age group. Error bars represent standard errors. The red line represents a linear regression model. **(E)** Distribution of oocyte sizes in the ovary throughout aging. (week 5 = 20 ovaries, week 10 = 15 ovaries, week 22 = 16 ovaries, week 31 = 18 ovaries, week 40 = 12 ovaries, week 50 = 10 ovaries, week 60 = 10 ovaries). The shaded region represents standard deviation. Insets show a magnified view of the 40-100 µm range.

Intriguingly, however, the fraction of transitioning oocytes — those exhibiting both squamous and cuboidal granulosa cell morphologies — remained stable at approximately 14% of the ovarian reserve across all ages, even when the total reserve declined by more than an order of magnitude (Fig. 2C–D). This suggests that, throughout the entire reproductive lifespan of the animal, and despite the large variation in oocyte numbers between isogenic individuals, the ovary maintains a proportionate number of oocytes primed for activation at any given time.

To determine whether external reproductive activity impacts oocyte depletion, we asked whether suppression of ovulation could slow the loss of the ovarian reserve. Ovulation suppression has been shown to reduce chromosomal mis-segregation in aging oocytes, thereby improving their quality and delaying functional decline^22^. However, its effect on the quantity of remaining oocytes has not been clearly established. To test this, we compared the ovarian reserve in females subjected to repeated mating with that of age-matched virgin controls. Mating began at 7 weeks and continued for 18 weeks, with animals undergoing multiple successful pregnancies. At 31 weeks, we quantified total oocyte numbers, along with primordial, transitioning, and growing follicles.

We observed no significant differences between mated and virgin females in any of these categories (Fig. S4E). These findings suggest that suppression of ovulation, while beneficial for oocyte quality, does not slow the rate of oocyte loss with age.

Together, these findings suggest that follicle recruitment is regulated at the tissue level and operates independently of ovulatory activity.

### The Ovarian Reserve Exhibits a Bimodal Size Distribution of Oocytes

Once activated, oocytes undergo dramatic growth, increasing their volume by approximately 64-fold^23^. How this growth is regulated remains poorly understood. If oocytes were activated at a constant pace and grew at a uniform rate, we would expect their sizes within the ovary to form a smooth continuum extending up to the ovulation stage. In contrast, if activation occurs in pulses or if growth is not continuous, we would anticipate depletions or accumulations at particular size ranges, reflecting pauses. To test these possibilities, we leveraged our quantitative 3D datasets across multiple ages and mapped the size distribution of all oocytes within intact ovaries.

We found that the size distribution of oocytes is bimodal: a dominant peak representing dormant oocytes, and a smaller secondary peak at approximately 60 µm (Fig 1G and Fig. 2E). This pattern suggests that oocytes encounter a developmental bottleneck or checkpoint at around 60 µm. The 60 µm size aligns with the late secondary–early antral stage, coinciding with the stage when follicles become gonadotropin-responsive^24^, raising the possibility of oocyte growth rates changing (i.e., faster growth after gaining gonadotropin sensitivity) during oogenesis. Indeed, recent research suggested a burst of oocyte growth right after this stage driven by a novel oocyte-specific organelle^25^.

Remarkably, the bimodal distribution is preserved throughout the reproductive lifespan of the animal, even though the ovarian reserve diminishes by more than an order of magnitude (Fig. 2E). This persistence indicates that the mechanisms governing the ∼60 µm bottleneck are robust to aging and independent of total oocyte number. Together, these findings suggest that oocytes experience a bottleneck when follicles become responsive to the gonadotropins, delaying or limiting further growth.

### Spatial Patterning of Oocytes Reveals Lateral Enrichment Along the Ovarian Axis

Oocyte distribution within the ovary is known to follow spatial cues^26^, but its full three-dimensional organization remains poorly characterized. For example, primordial oocytes tend to localize to the cortical region, while growing oocytes are typically found closer to the medulla. To better characterize these spatial patterns, we leveraged the full 3D positional data from our whole-ovary imaging pipeline to quantify oocyte organization across the entire ovarian volume.

We began by generating isosurface renderings of each ovary, revealing a consistent, stereotypical bean-like morphology. This visualization clearly delineated the concave medial side of the ovary, known as the *hilum* — the entry point for nerves and blood vessels — which served as a reproducible anatomical landmark across samples. To systematically quantify radial organization, we developed a spatial framework based on the ovary’s geometry. We defined the long and short axes of each ovary and fit a spline along the central longitudinal axis to serve as an internal reference line, or “ruler,” from which the position and angle of each oocyte could be measured. This allowed us to assess distribution patterns in a normalized coordinate space, enabling comparison across animals and age groups (Fig. 3A).

**Figure 3.**
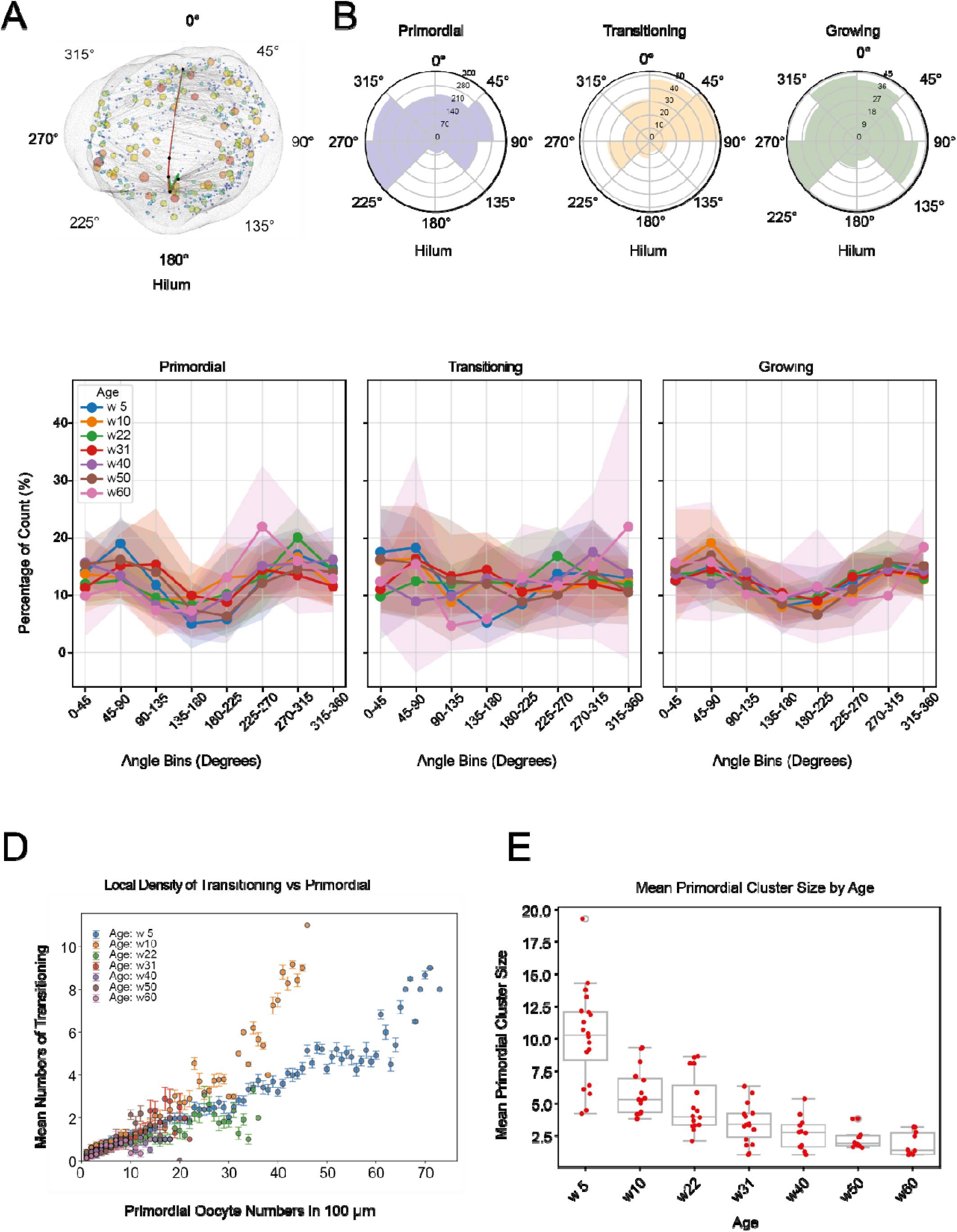
Regions of high primordial oocyte density show higher activation. **(A)** Iso-surface rendering of a 5-week-old C57BL/6J ovary in light grey. Segmented oocytes are plotted as spheres and color-coded by their diameter. The red solid line represents the spline curve used for spatial reference. 0° corresponds to the superior side of the ovary, 45– 135° and 225–315° represent the medio-lateral regions, and 180° corresponds to the hilum. **(B)** Quantification of the radial distribution of oocytes shown in (A), plotted separately for primordial (left), transitioning (middle), and growing (right) oocytes. **(C)** Quantification of the radial distribution of oocytes during aging, each solid line represents the mean value and shaded area represents the standard deviation (week 5 = 8 ovaries, week 10 = 5 ovaries, week 22 = 6 ovaries, week 31 = 9 ovaries, week 40 = 5 ovaries, week 50 = 7 ovaries, week 60 = 5 ovaries). Percentages of oocytes contained in bins of 45° are shown for primordial oocytes (left), transitioning oocytes (middle) and growing oocytes (right). **(D)** Quantification of the number of transitioning oocytes as a function of primordial oocyte cluster size. Primordial cluster size is defined as the number of primordial oocytes within a radius of 100 µm from each primordial oocyte. Each dot represents the mean number of transitioning oocytes associated with a given cluster size, color-coded by age. Error bars indicate standard deviations. (week 5 = 20 ovaries, week 10 = 15 ovaries, week 22 = 16 ovaries, week 31 = 18 ovaries, week 40 = 12 ovaries, week 50 = 10 ovaries, week 60 = 10 ovaries). **(E)** Quantification of the average primordial cluster size. Each dot represents the average primordial cluster size per ovary across different ages.

Our analysis revealed that primordial oocytes were consistently depleted from both the concave (hilum) and convex poles of the ovary, while showing consistent enrichment along the lateral mid-regions of the long axis. This pattern was not limited to dormant oocytes: transitioning and growing oocytes exhibited the same spatial bias, and this distribution was maintained throughout aging (Fig. 3B–C). These findings align with recent observations showing that the ovarian surface epithelium is sparsely populated near the hilum^27^.

While follicle activation has been proposed to occur stochastically^28,29^, local microenvironmental factors can influence whether a primordial oocyte gets activated or remains dormant^30,31^.

To test whether neighboring-follicle density influences oocyte activation, we performed a spatial density analysis of primordial oocytes across the ovarian volume. For each primordial oocyte, we quantified the number of neighboring primordial oocytes within a 100 µm radius. We observed highly heterogeneous local densities of follicles, which decrease with age: at 5 weeks, any primordial oocyte has, on average, over 10 neighbors, whereas at 60 weeks, this number is reduced to approximately 2.5 (Fig. 3E). We did not observe persistent high-density clusters of primordial follicles at later ages, suggesting that the high local density of primordial follicles does not inhibit their activation. Instead, we found that higher local primordial oocyte densities were associated with increased numbers of transitioning oocytes (Fig. 3D), irrespective of age. To further check this correlation, we also examined the inverse scenario: whether a high local density of transitioning oocytes was associated with increased numbers of primordial oocytes. We found a positive correlation in this case as well (Fig. S5A). These results suggest that a high density of primordial oocytes in a given region is not inhibitory towards follicle activation, as previously suggested for prepubertal mouse ovaries^26^. On the contrary, we find that more primordial oocytes in a given region correlate with more active oocytes in the same area.

### Mathematical Modeling Captures the Kinetics of Ovarian Reserve Depletion During Aging

Current technologies do not permit in vivo longitudinal studies of the ovary; therefore, we leveraged our large, age-spanning dataset to model the dynamics of ovarian reserve depletion and infer the underlying regulatory processes. To construct this model, we classified oocytes into four developmental categories based on follicular morphology and size: primordial (diameter ≤ 20 µm), primary (20 µm < diameter ≤ 40 µm), secondary (40 µm < diameter ≤ 60 µm), and antral (diameter > 60 µm). These four stages were chosen because they represent major morphological milestones in folliculogenesis^32^.

This classification enabled us to plot stage-specific oocyte distributions across all time points, effectively capturing how the size profile of the ovarian reserve changes with age (Fig. 4A). To account for natural variation in ovarian size, we normalized oocyte counts to the total organ volume (Fig. S4C–D), which reduced the spread of the distributions and removed outliers (Fig. S5B).

**Figure 4.**
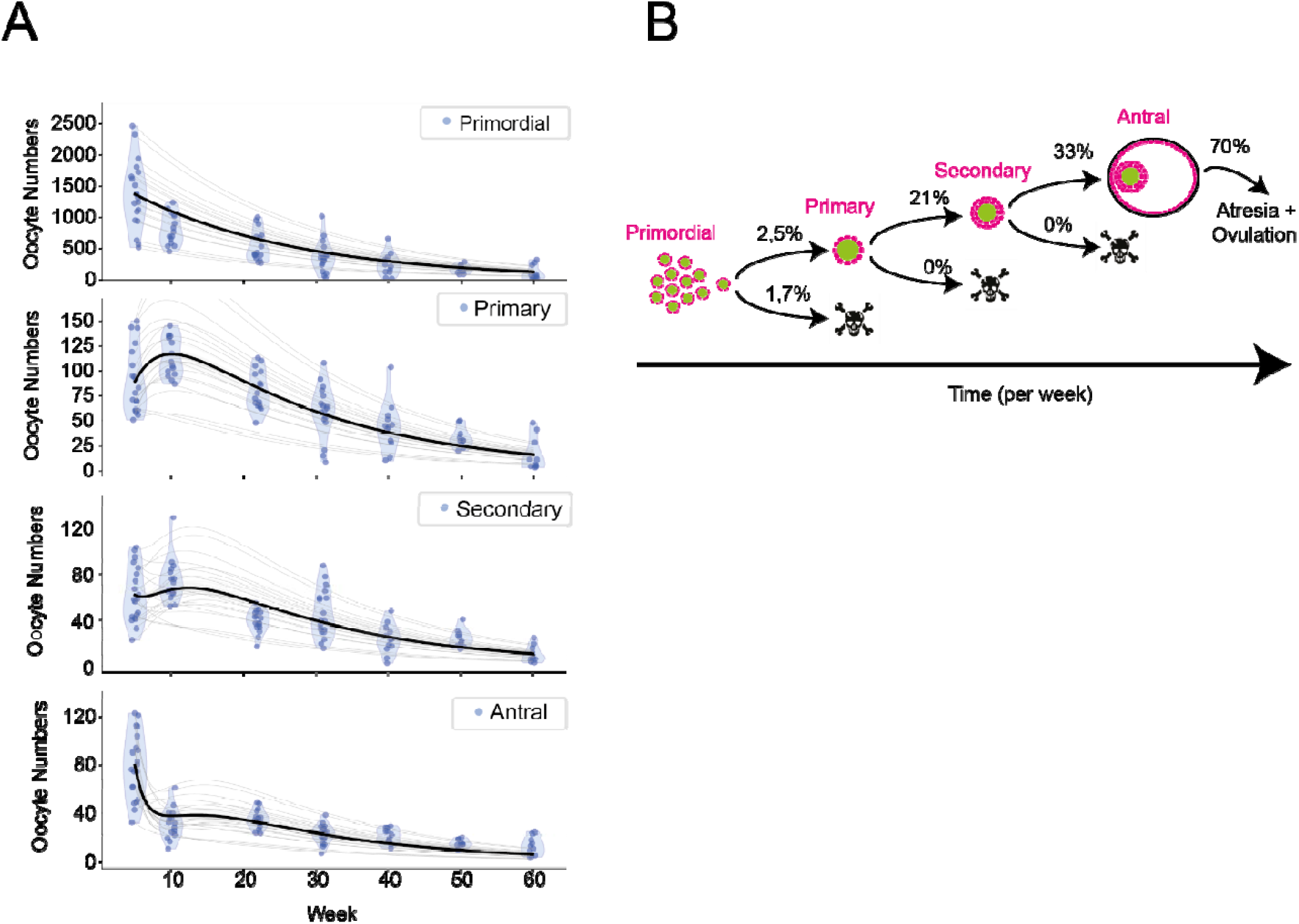
A mathematical model recapitulates ovarian aging dynamics. **(A)** Violin plots showing the number of oocytes per ovary across different age groups, categorized by oocyte stage: primordial, primary, secondary, and antral. Each dot represents an individual ovary. Light grey lines indicate model fits based on varying initial conditions, while the black line represents the fit starting from the mean values per each category of week 5 animal. **(B)** Schematic representation of the mathematical model. The model suggests that oocyte death is minimal during the primordial, primary, and secondary stages, but increases significantly at the antral stage. An activation rate of 2.5% of oocytes per week is sufficient to maintain the follicular pool for at least 60 weeks

Oocyte activation is widely accepted as a unidirectional process^6^. Once an oocyte initiates growth, it either continues maturing or undergoes atresia, but it does not revert to dormancy.

Conceptually, this process can be represented as a sequence of transitions: primordial oocytes either progress to the primary stage or die; primary oocytes either advance to the secondary stage or die; secondary oocytes either develop into antral follicles or die; and antral follicles either ovulate or degenerate.

We modeled this developmental trajectory over time using a system of ordinary differential equations (ODEs) to describe the rates of change for each transition. In this framework, follicle progression from primordial to antral stages (denoted *P*_*i*_) is governed by stage-specific transition rates (*k*_*i*_) and death rates (*d*_*i*_), where index *i* represents each step in the sequence.

The parameters of these equations were fitted to experimental observations of oocyte numbers in each subgroup (Fig. 4A), yielding the following results:

k_1_ (primordial → primary) = 0.025808 ± 0.005288 week^−1^

k_2_ (primary → secondary) = 0.217252 ± 0.040008 week^−1^

k_3_ (secondary → antral) = 0.337478 ± 0.056654 week^−1^ d_1_ (primordial death) = 0.017856 ± 0.007404week^−1^

d_2_ (primary death) = 0.0000 ± 0.0000 week^−1^

d_3_ (secondary death) = 0.0000 ± 0.0000 week^−1^

d_4_ (antral death and ovulation) = 0.722362 ± 0.147043 week^−1^

All constants are expressed in units of week □ ^1^. These values indicate that, each week, approximately 2.5% of primordial oocytes are activated, 21.7% of primaries transition to the secondary stage, and 33.71 % of secondaries develop into antral follicles. These results provide the first quantitative estimates of the oocyte activation and growth dynamics in the mammalian ovary (Fig. 4A–B). Our model predicts very low death rates for primary and secondary oocytes in adult ovaries and suggests that most oocyte loss occurs at the antral stage, consistent with findings in the literature^20,33^ and further supporting our model’s validity.

Given the high inter-individual variability in the ovarian reserve size, we next tested whether the model could solve the equations starting from the initial conditions of each individual 5-week-old ovary. Notably, the model remained stable across all these starting conditions (Fig. 4A).

Taken together, these analyses demonstrate that a mathematical model can recapitulate the kinetics of ovarian reserve depletion across the mouse reproductive lifespan. This framework provides the first quantitative estimates of follicle recruitment, growth, and death rates, and establishes a basis for further refinement and hypothesis generation.

## Discussion

The duration of a female’s fertility is determined by the initial number of her oocytes and the rate at which they are exhausted^8^. Therefore, understanding the kinetics of the ovarian reserve is key to understanding ovarian aging. Here, we provide a high-resolution view of ovarian reserve dynamics across the reproductive lifespan of a mammal using whole-organ 3D imaging, AI-driven segmentation, and stage-specific classification of oocytes. By imaging intact mouse ovaries and quantifying over 100,000 oocytes across ages, we first uncovered striking inter-individual variability in oocyte numbers among genetically identical mice, an aspect that has rarely been appreciated due to the lack of quantitative, whole-organ analyses. This variability becomes evident only through comprehensive 3D quantification of ovaries of several animals, enabling us to uncover age-dependent trends that were previously inaccessible. Notably, while the total ovarian reserve declines dramatically with age, the proportion of transitioning follicles remains stable, suggesting a homeostatic mechanism that regulates follicle activation. Our results support the hypothesis that oocyte activation can be sensed at the tissue level and suggest that the ovary possesses a mechanism to monitor the total number of oocytes across a wide distribution of ovarian reserve sizes and animal ages.

Quantifying the ovarian reserve in intact ovaries has been challenging, as high magnification is required to identify primordial oocytes—typically around 15 µm in diameter—while still maintaining a field of view large enough to accommodate millimeter-scale samples. Two recent studies combined tissue clearing with high-resolution microscopy and machine learning approaches to automate the quantification of oocytes^34,35^. Our method is a major improvement on these approaches, as it is sufficiently generalizable to work on adult mouse ovaries across the reproductive lifespan as well as on human ovaries and enables segmentation of whole oocytes, including their cytoplasm, thereby allowing both volumetric analyses and lifespan-scale quantification within a single framework.

A central unresolved question in ovarian biology is why some oocytes become activated while the majority remain dormant at any given time. Our data provide new insights into this process, revealing that regions with a high density of primordial oocytes contain a greater number of recently activated oocytes compared to less dense regions. This observation challenges the long-standing “high-density inhibition” hypothesis in mouse ovaries^26^.

We also found that the size distribution of all oocytes within the ovary is bimodal, with a first peak at ∼15 µm representing the pool of primordial oocytes, and a second peak at ∼60 µm corresponding to the late secondary–early antral stage (Fig. 2E). This suggests the existence of a developmental bottleneck or checkpoint around 60 µm, at which many oocytes may pause, possibly awaiting additional signals from granulosa cells, hormones, or metabolic cues before progressing further. This is an important stage of folliculogenesis, as new regulatory factors come into play. After this stage, continued follicle growth depends on gonadotropic hormones such as follicle-stimulating hormone (FSH) and luteinizing hormone (LH) that promote granulosa/cumulus cell proliferation, survival, and estradiol production, all of which are essential for further follicle development. Further investigation into the signals controlling this bottleneck will be essential to fully understand oocyte growth dynamics.

Finally, the scale and resolution of our dataset allowed us to explore the kinetics of ovarian reserve depletion through mathematical modeling. In building this model, we assumed that oocyte growth and death rates remain broadly similar across aging ovaries. We acknowledge that this is an assumption that may require refinement as additional data become available. The model predicted minimal cell death at the primordial, primary, and secondary stages, with the majority of oocyte loss occurring at the antral stage. This prediction is consistent with previous experimental reports^20,33^, thereby reinforcing the robustness of our model’s assumptions. We believe this framework establishes a foundation for generating new hypotheses and refining future models of ovarian reserve dynamics.

Altogether, by merging whole-organ imaging with quantitative modeling, our work reframes the ovarian reserve as an actively regulated system rather than a passive reservoir, opening new avenues for understanding the principles governing female reproductive longevity and their broader implications for tissue maintenance and aging.

## Methods

### Whole-mount ovary immunostaining

Ovaries were obtained from C57BL/6J mice aged 5–60 weeks. After dissection, the bursa was manually removed, and ovaries were fixed for 1 h at room temperature (RT) in 2% PFA in PBS, then transferred to 80% methanol/DMSO at −20 °C for at least one overnight, as previously described^34^. Ovaries were then washed in 50% methanol/PBS, 25% methanol/PBS, and PBS for at least 30 min each step and permeabilized for 5 days at RT with gentle agitation. Ovaries were blocked for 1 day at RT in 10% normal goat serum (NGS), 1% Triton X-100, and 0.05% NaN3, then incubated with primary antibody for at least 3 days, washed 4 × 30 min in 1% Triton X-100/PBS, incubated with Alexa-conjugated secondary antibody for at least 3 days at RT, and washed again 4 × 30 min in 1% Triton X-100/PBS.

### Human ovary cortex immunostaining

Ethical committee permission to use human ovary samples was obtained from the Comité Étic d’Investigació Clínica CEIC-Parc de Salut MAR (Barcelona) and Comité Ético de Investigación Clínica–Hospital Clínic de Barcelona with approval number HCB/2023/0777. Written informed consent was obtained from both participants.

Pieces of the human ovarian cortex were donated by two individuals aged 25 years. Tissue was fixed in 4% PFA for 4 h at room temperature (RT), then transferred to 80% methanol/DMSO at −20 °C for at least one overnight. Samples were permeabilized for ≥8 days in 2% Triton X-100/PBS, blocked overnight in 10% NGS, 1% Triton X-100, and 0.05% NaN3, incubated for ≥4 days with primary antibody, and then incubated for ≥4 days with Alexa-conjugated secondary antibody.

### Mounting ovaries in agarose block and BABB clearing

The mouse ovaries or human ovarian cortex pieces were washed twice in Milli-Q water (to remove PBS salts) and transferred to a 2 mL Eppendorf tube containing 2 mL of 1% low-melting agarose. The ovary was positioned in the middle of the tube, and the tube was laid on its side on ice to accelerate solidification. After ∼30 min, the agarose had solidified; the end of the tube was cut with a blade and the agarose cylinder containing the ovary was extracted. The cylinder was further trimmed to fit the SPIM sample holder. Samples were then transferred to a 20 mL glass tube and incubated in 100% methanol for at least one (O/N); methanol was subsequently removed and tissue clearing was performed in benzyl alcohol:benzyl benzoate (BABB; 2:1, vol/vol)

### SPIM Microscopy

Whole ovaries were labeled with an antibody against DDX4 and the dye TO-PRO-3, to label oocytes and DNA, respectively. The ovaries were imaged using a Luxendo MuViSPIM equipped with a 10x detection lens and two-side illumination set-up, resulting in a resolution of a pixel size of 0.867 μm. Images were spaced 2.5 μm in Z. Excitation at 488 nm was used for detecting DDX4 positive oocytes and 642 nm for detecting TO-PRO-3 labelled nuclei. In order to cover the whole volume multiple tiles were acquired. Stitching of the tiles were performed either using MosaicExplorerJ^36^ or using the Luxendo image software.

### Confocal microscopy

Whole ovaries labeled with an antibody against DDX4 to label oocytes and TO-PRO-3 dye to label DNA and were imaged using an SP8Stellaris system equipped with a 63X glycerol lens. We used the function tile-scan in order to cover a large ovary area.

### Oocyte recognition using BiaPy

To specifically address the task of oocyte segmentation, i.e instance segmentation task, we employ a bottom-up approach to train the network to predict three channels: binary foreground mask (F) and instance contour map (C), and an Euclidean distance map to the center of the instance (Dc). This model can be called “FCDc”, following the nomenclature and the successful combination of these channels in other state-of-the-art methods^37-42^. After prediction, all outputs are thresholded and combined to enhance object delineation. A connected components algorithm is then applied to the combined mask to identify discrete, non-overlapping oocyte instance seeds. These seeds are subsequently used as markers in a marker-controlled watershed algorithm^43^, which incorporates three key components: (1) the inverted foreground probability map as the input image (defining the topographic surface), (2) the instance seeds as the marker image (indicating the initiation points of the flooding process), and (3) a binarized version of the foreground mask (B) as the constraint mask (limiting the extent of region expansion). This integrated pipeline enables the effective segmentation of individual oocyte instances. A visual representation of the process is provided in Fig. S1A.

The model has five-levels with 48 filters in the initial level that get summed by 16 on each level (until 96 in the bottleneck), ELU activation functions and transposed convolutions to perform the upsampling in the decoder. We used batch normalization for regularizing the model. The model was trained using a patch size of 40×128×128 and a kernel-size of 3 in all convolutions.

Our approach is fully integrated within the BiaPy library^16^, using Pytorch version 2.4. As data augmentation techniques we employed horizontal and vertical flips, elastic transformations and gridmarks^44^. We set aside 10% of the training samples for validation. Cross entropy was used as a loss function, with a learning rate of 1e-4, employing a cosine-decay scheduler with warm-up^45^, and ADAMW optimizer^46^. The model was trained until convergence using eight NVIDIA GeForce RTX3090.

### Data Analysis

#### Ovarian reserve quantification

After visual inspection and correction of the segmented mask, we calculated the 2D bounding box around each oocyte using MorpholibJ Fiji plugin^47^. From the 2D bounding box we can obtain the diameter calculating the square root of the product of the short side by the long side of the bounding box:√(a·b). For choosing the threshold between primordial and transitioning oocytes we use the mean +/− the st. deviation of the values measured in Fig. S2D. We got values from 16,30 μm to 20,11 μm in diameter. Oocytes with diameter from 10 to 16,30 μm were classified as primordial; 16,30 to 20,11 as transitioning and oocytes greater than 20,11 μm were classified as growing.

#### Primordial peak alignment

To account for slight differences (1-2 microns) in oocyte sizes across ovaries, we aligned all individual primordial peaks to the average primordial peak of the entire dataset. First, we calculated the average primordial peak across all ovaries, and then aligned each individual primordial peak to this reference.

#### Oocyte radial quantification

Iso-surface rendering of the ovarian surface was performed from the raw image stacks using the Vedo library^48^ to visualize the three-dimensional morphology of the ovary. From the corresponding segmented masks, the center of mass of each oocyte was computed using the MorphoLibJ plugin in Fiji. A spline was then manually positioned through the ovary, connecting the center of the short axis and extending along the long axis. Due to the geometry of the ovary, this midline typically lies within a single plane, allowing a consistent definition of its orientation. Based on this reference, the angular distribution of oocytes was measured, distinguishing dorsal oocytes (located at small angles, facing the concavity of the spline) from ventral oocytes (located at angles approaching 180°).

#### Modelling the ovarian reserve over aging

The oocyte size distribution was partitioned into four diameter-based classes: **primordial** (10– 20 µm), **primary** (>20–40 µm), **secondary** (>40–60 µm), and **antral** (>60 µm). Class boundaries were defined to be non-overlapping. A set of differential equations is then solved using as initial conditions the number of oocytes present in each category at week 5. The minimization is then performed using the Sequential Least Squares Programming (SLSQP) algorithm as implemented in the *SciPy* library.

During fitting to the experimental oocyte counts, the least-squares constrained minimization leads to zero values for the death rates for primary and secondary type oocytes (*d*_*2*_ and *d*_*3*_). As this is consistent with what is known in the literature^20,33^, these two parameters were fixed to zero to obtain our final results (Fig. 4A).

## Acknowledgments

We would like to thank the EMBL’s Mesoscopic Imaging Facility and CRG’s ALMU for the support and assistance in this work. We are also grateful to the PRBB Animal Facility for assistance with animal care. We are grateful to Julien Colombelli and Sebastian Tosi (IRB-Barcelona) for helping with the image stitching. We thank Alba Granados (IsardSAT-Barcelona) and Lucas Rapez for helping with the python scripts. We are grateful to Gabriele Zaffagnini (CECAD-Cologne) and Nicholas Stroustrup (CRG-Barcelona) for their comments on the manuscript.

We acknowledge the support of the Spanish Ministry of Science and Innovation through the Centro de Excelencia Severo Ochoa (CEX2020-001049-S, MCIN/AEI/10.13039/501100011033) and the Generalitat de Catalunya through the CERCA program. M.M. and J.S. were supported by funds from EMBL. This work was partially supported by grant GIU23/022, funded by the University of the Basque Country (UPV/EHU), and grant PID2021-126701OB-I00, funded by the Ministerio de Ciencia, Innovación y Universidades, MICIU/AEI /10.13039/501100011033/, and “ERDF A way of making Europe”. E.B. acknowledges funding from the European Research Council Consolidator Grant (ACTIVEDORMANCY_ 101088824).

