## supplementary for "3D Mapping of Intact Ovaries Reveals the Aging Dynamics of the Ovarian Reserve"

Figure_S1

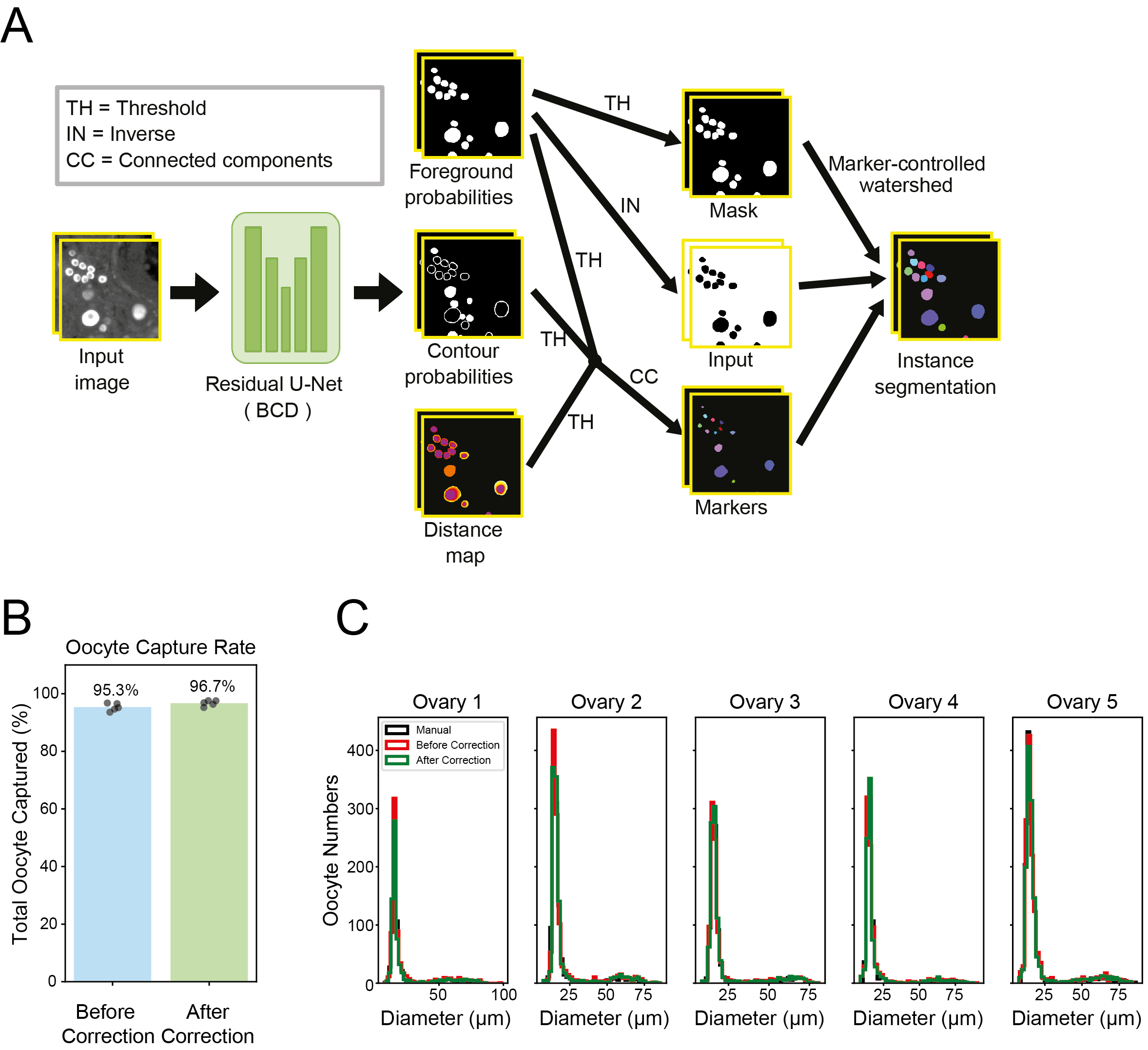

Figure S1. **Validation of the automated oocyte detection and segmentation pipeline.**

**(A)** Schematic representation of the oocyte recognition pipeline. Model outputs, e.g., foreground, contours and distances-to-the-center, are integrated and used as input for a marker-controlled watershed algorithm **(B)** Comparison of the total number of oocytes counted manually versus automatically, before and after user correction. Each dot represents an ovary from a 5-week-old C57BL/6J mouse. **(C)** Histogram showing the full oocyte size distribution obtained from manual segmentation, automatic segmentation before correction, and after user correction, for each ovary shown in (B).

Figure_S2

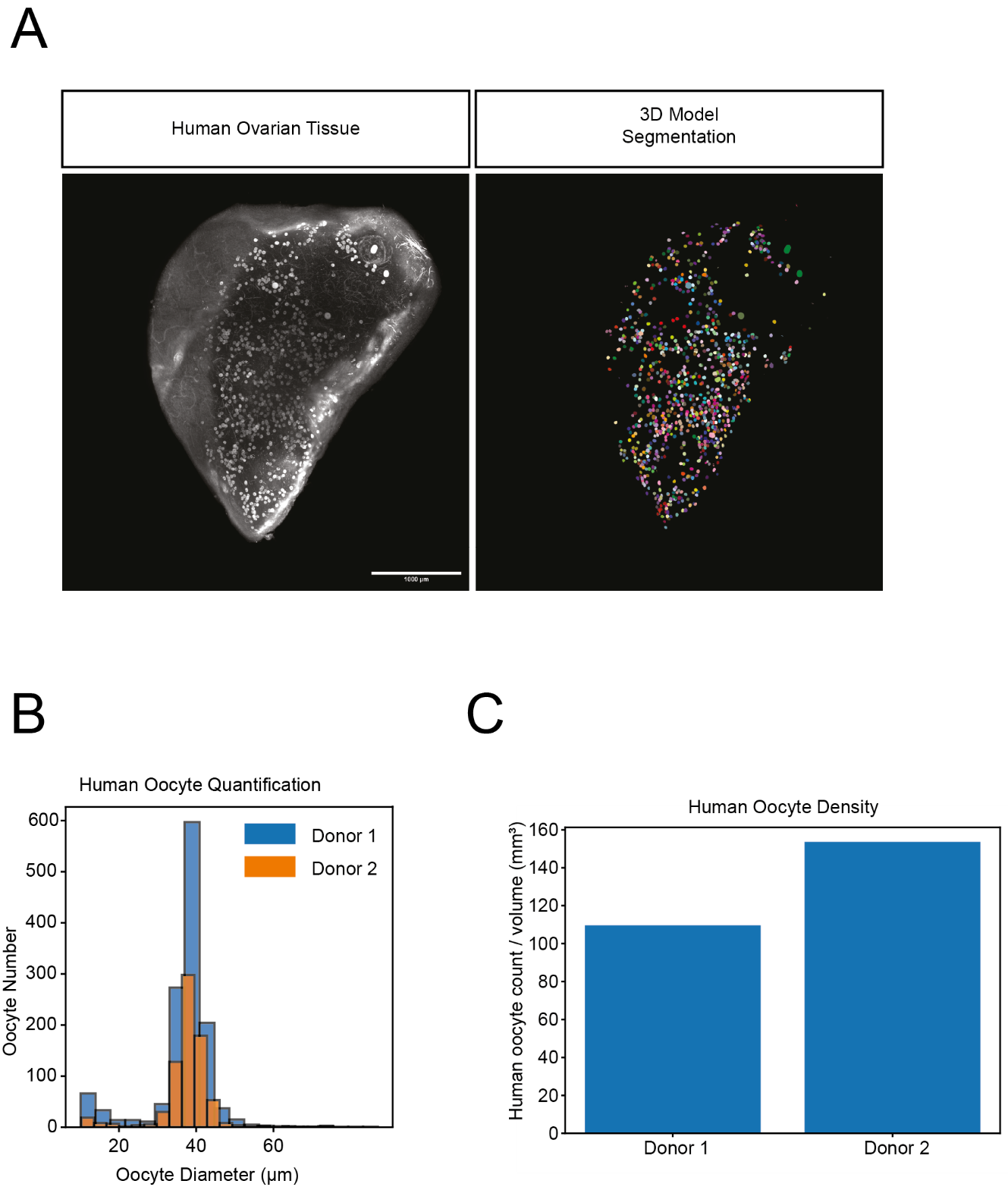

Figure S2. **Automated quantification of oocytes in cortical strips of human ovarian tissue**

**(A)** SPIM image of a fragment of human ovarian cortex from a 25-year-old individual labeled with an antibody against the oocyte marker DDX4 (left), and the corresponding automated segmentation of all oocytes (right). Scale bar: 1 mm. **(B)** Quantification of total oocyte numbers from two ovarian cortex fragments obtained from two different individuals. **(C)** Bar plot showing oocyte count per unit volume (mm³) for two donors. Oocyte counts were obtained from the segmented dataset shown in (B) and normalized to each sample’s measured volume.

Figure_S3

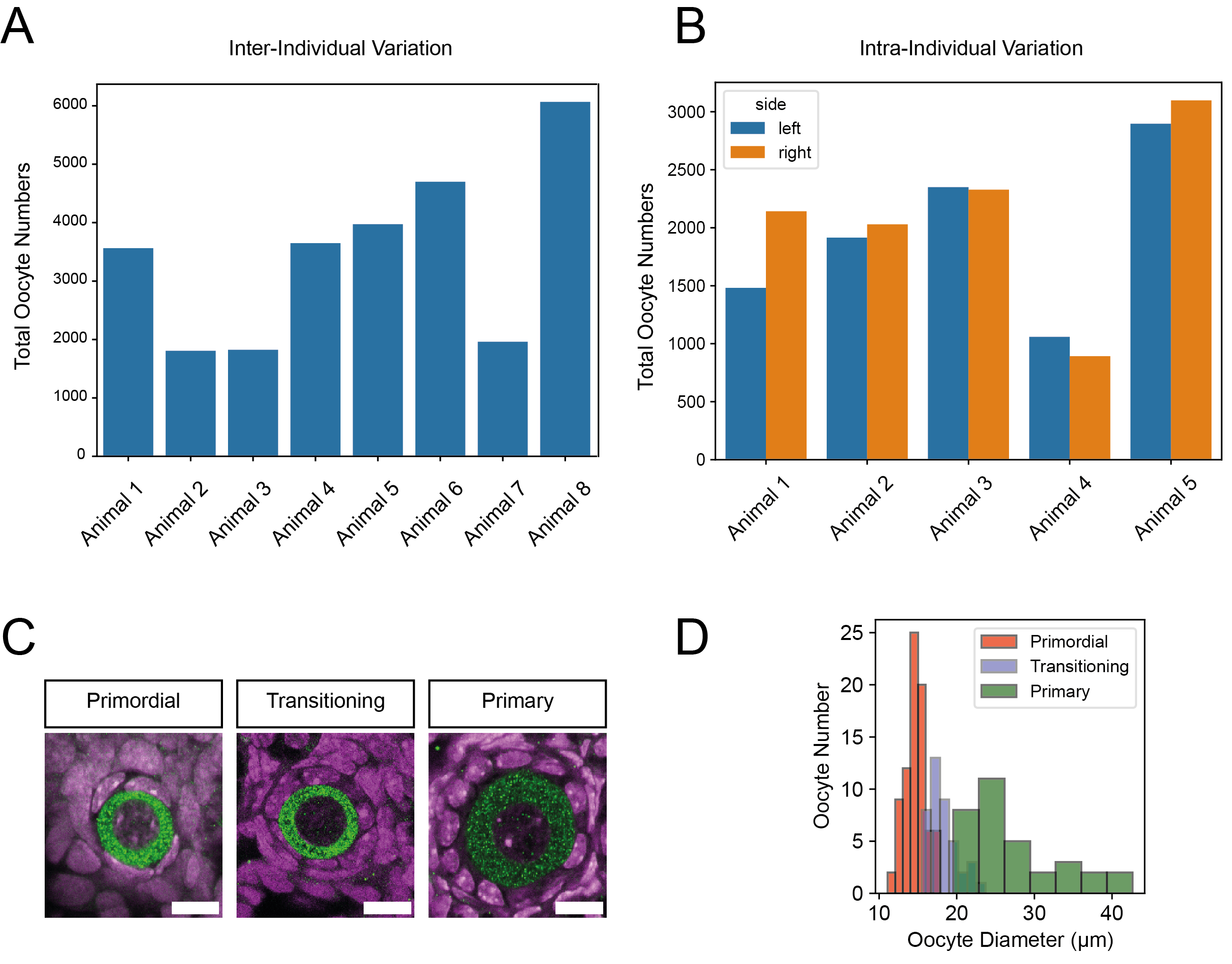

Figure S3. **Inter- and intra-individual variation in the ovarian reserve**

**(A)** Inter-individual variation in 5-week-old C57BL/6J inbred mouse ovaries. Each bar represents the combined oocyte count from both ovaries of an animal. **(B)** Intra-individual variation in the ovarian reserve in 5-week-old C57BL/6J inbred mice, showing the oocyte counts for left and right ovaries separately. **(C)** Representative confocal images of primordial, transitioning, and primary oocytes. Oocytes are labeled in green (anti-DDX4), and nuclei are labeled in magenta (TO-PRO-3). Scale bar is 10 μm. **(D)** Quantification of oocyte diameters for primordial, transitioning, and primary oocytes.

Figure_S4

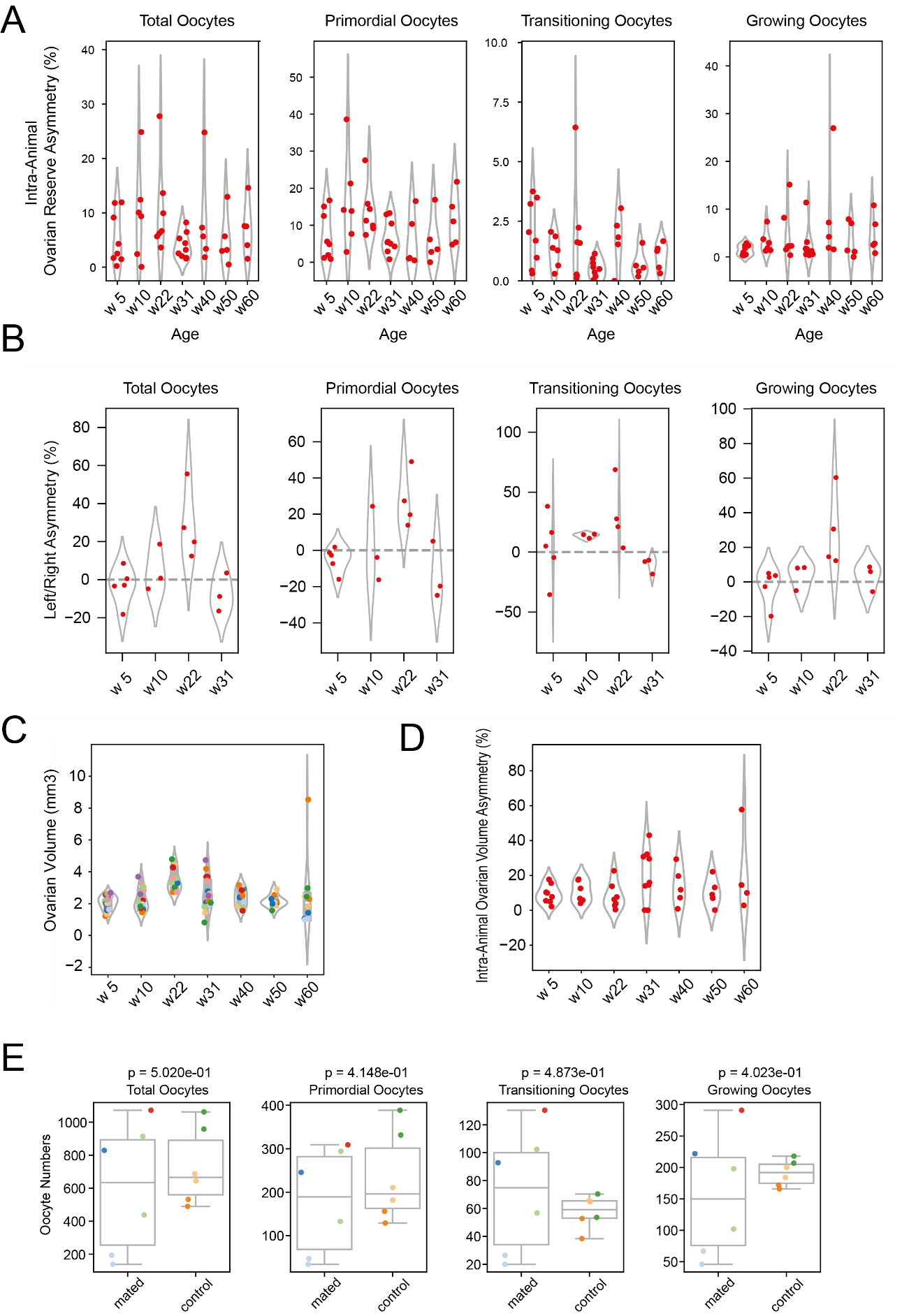

Figure S4. **Intra-individual variation and ovulation-dependent changes in the ovarian reserve across aging.**

**(A)** Intra-individual variation in ovarian reserve across different ages. Each dot represents the percentage difference in total, primordial, transitioning and growing oocyte number between the ovary with the higher reserve and the one with the lower reserve from the same animal. **(B)** Left–right asymmetry in ovarian reserve across different ages. Each dot represents the percentage difference in total, primordial, transitioning, and growing oocyte numbers between the left and right ovary of the same animal. **(C)** Quantification of ovarian volume during aging. Each dot represents an ovary. **(D)** Intra-individual variation in ovarian volume across different ages. Each dot represents the percentage difference in ovarian volume between the larger and smaller ovary of the same animal. **(E)** Quantification of total, primordial, transitioning, and growing oocytes in virgin versus mated animals that successfully delivered live offspring. All the animals were sacrificed at 31 weeks of age. Each dot represents one ovary. Statistical comparisons between the two conditions within each category were performed using an unpaired two-sample t-test assuming unequal variance.

s

Figure_S5

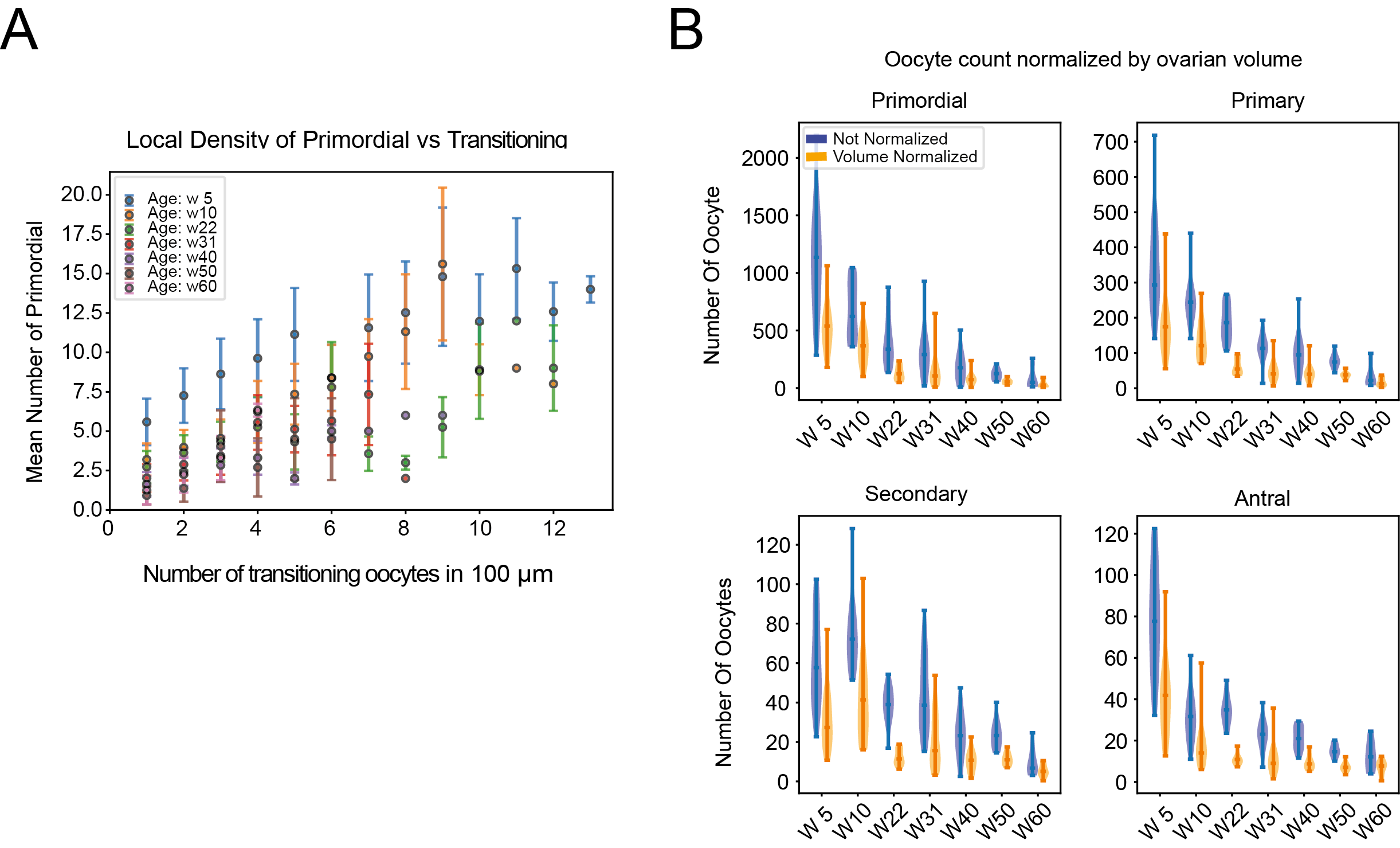

Figure S5. **Local density relationships and volume normalization of oocyte counts across aging.**

**(A)** Quantification of the number of primordial oocytes as a function of transitioning oocyte cluster size. Transitioning [oocyte] cluster size is defined as the number of transitioning oocytes contained in a distance of 100 µm from each transitioning oocyte. Each dot represents the mean number of primordial oocytes associated with a given cluster size, color-coded by age. Error bars indicate standard deviations. (week 5 = 20 ovaries, week 10 = 15 ovaries, week 22 = 16 ovaries, week 31 = 18 ovaries, week 40 = 12 ovaries, week 50 = 10 ovaries, week 60 = 10 ovaries). **(B)** Violin plots showing the number of oocytes per ovary across different age groups, grouped by stage: primordial, primary, secondary, and antral. Orange violins depict raw oocyte counts, while blue violins represent data normalized by ovarian volume. Normalization reduces the spread of the data, suggesting that ovarian volume contributes to the variability in oocyte numbers. Each dot represents an individual ovary.

Movie S1. **Whole-mount ovary staining.**

SPIM z-stack of a five-week-old C57BL/6J ovary, labeled with anti-DDX4 (green) and TO-PRO-3 (magenta)

Movie S2. **Whole-mount ovary staining and oocyte detection.**

SPIM z-stack and 3D reconstruction of a five-week-old C57BL/6J ovary labeled with anti-DDX4, overlaid with the corresponding segmentation mask.

Movie S3. **Imaging and** **quantification of oocytes in cortical strips of human ovarian tissue.**

SPIM z-stack of a strip of human cortex labeled with anti-DDX4 and overlaid with the corresponding segmentation mask.
